# Proteome landscape of B-cell malignancies identifies mantle cell lymphoma protein signature

**DOI:** 10.64898/2026.03.02.709116

**Authors:** Samantha A. Swenson, C. Bea Winship, Kasidy K. Dobish, Karli J. Wittorf, Henry Chun Hin Law, Julie M. Vose, Timothy Greiner, Michael R. Green, Nicholas T. Woods, Shannon M. Buckley

**Affiliations:** Department of Biochemistry and Molecular Biology, University of Nebraska Medical Center, Omaha, NE, USA; Department of Genetics, Cell Biology and Anatomy, University of Nebraska Medical Center, Omaha, NE, USA; Fred and Pamela Buffett Cancer Center, University of Nebraska Medical Center, Omaha, NE; Department of Internal Medicine, Division of Hematology & Hematopoietic Malignancies, University of Utah, Salt Lake City, UT USA; Huntsman Cancer Institute, University of Utah, Salt Lake City, UT USA; Creighton University School of Medicine, Omaha, NE, USA; Eppley Institute, University of Nebraska Medical Center, Omaha, NE, USA; Department of Pathology and Microbiology, University of Nebraska Medical Center, Omaha, NE, USA; Department of Lymphoma and Myeloma, Division of Cancer Medicine, The University of Texas MD Anderson Cancer Center, Houston, TX, USA; Department of Genomic Medicine, University of Texas MD Anderson Cancer Center, Houston, TX, USA; Center for Cancer Epigenetics, University of Texas MD Anderson Cancer Center, Houston, TX, USA

## Abstract

Mantle cell lymphoma (MCL) is one of the deadliest forms of Non-Hodgkin’s B-cell lymphoma. Typically, patients present with both overexpression of CyclinD1 and secondary mutations identified by genomic sequencing. Although MCL patients may initially respond to treatment, they eventually relapse and succumb to disease, highlighting the essential need to identify new targets for treatment. Here we performed proteomic profiling of healthy B cells and three different forms of B-cell malignancies, including MCL, to define the proteomic signature of MCL. We compared the proteome of each to MCL and identified 10 proteins that are specifically upregulated in MCL. Of these 10 proteins, seven of them show no transcriptional changes and have been overlooked by conventional RNA expression analysis. Further analysis of the proteomic signature reveals potential avenues for dual targeting in CAR T-cell therapy and provides guidance for personalized therapeutics based on protein expression.

**STATEMENT OF SIGNIFICANCE:** We present a resource defining the protein landscape of MCL, CLL, and FL as compared to healthy b cells identified utilizing quantitative proteomics from primary patient samples. Applied to MCL, our results identify 10 proteins specifically upregulated in MCL that may prove to be therapeutic targets to treat the disease.

## INTRODUCTION

Malignancies affecting the hematopoietic system make up approximately 8-10% of all human cancers.^1^ The two most common adult hematopoietic malignancies in western countries are chronic lymphocytic leukemia (CLL) and follicular lymphoma (FL). Both are B-cell malignancies with incidences of 25-30% of all adult leukemias and 20-40% of all Non-Hodgkin’s Lymphomas (NHL), respectively.^2^ Current therapies have improved treatment options and overall survival (OS). CLL has an ∼87% 5-year OS, in part due to the BTK inhibitor Ibrutinib being approved by the FDA in 2016 for frontline treatment.^3^ Likewise, FL has an ∼90% 5-year OS associated with the addition of the monoclonal antibody rituximab being used in conjunction with chemotherapeutics.^4^

Unfortunately, not all B-cell malignancies have similar survival rates. In the case of mantle cell lymphoma (MCL), an aggressive NHL that accounts for ∼4-6% percent of all lymphoma cases in the United States, patients have only an ∼40% 5-year survival.^2^ Besides the standard high dose chemotherapy regimen, limited treatments are available, resulting in poor outcomes.^2^ Recently, chimeric antigen receptor T-cell (CAR-T) therapy targeting the B-cell marker CD19 has been approved by the FDA for treatment in relapse/refractory MCL.^5^ Although this provides a new therapeutic option for MCL patients, it is not considered a universal cure and many patients still relapse.^5^ Additionally, the proteasome inhibitor Bortezomib was approved for MCL patients with relapse or refractory disease in 2013; however, discrepancies between survival of these patients show a significant need for new first-line therapeutics.^6^

This study aims to garner a better understanding of MCL pathogenesis and create a resource that can be used for the identification of potential novel therapeutic targets to improve patient prognosis. Currently, most research has focused on gene expression profiling (GEP) of patients to identify molecular predictors of disease; however, these studies are not extensively validated at the protein level where targeted therapies take effect. Additionally, current proteomic studies are heavily reliant on cell lines which cannot accurately mimic patient tumors.^7, 8^ When patient samples have been utilized, the comparison has only been made to healthy donor (HD) and not between disease types.^9,10^ The one instance where disease types have been compared (MCL to CLL), only transcriptomics was utilized.^11^ To address this gap in knowledge, we utilized 35 primary patient samples to identify an MCL-specific proteomic signature by comparing MCL to CLL, FL, and HD B cells by quantitative mass spectrometry. The minimal overlap observed between proteomic and GEP analyses emphasizes the importance of developing such a resource. Of particular interest are proteins we have identified as differentially expressed on the plasma membrane, as these may be ideal targets for CAR-T therapy. We also identified patient-specific variations in the protein abundance of proteasomal subunits suggesting that addition of Bortezomib to the treatment regimen may benefit some patients, but not others. Finally, our results identify 10 proteins specifically upregulated in MCL. Of these 10 proteins, seven are unchanged at the mRNA level, and would therefore not be identified by GEP. Importantly, three of these proteins have never been associated with any lymphoid malignancy, making them a novel finding.

## RESULTS

### Proteomics signatures of MCL, CLL, and FL

To identify the protein landscape of MCL, CLL, and FL, we performed quantitative tandem mass tag (TMT) mass spectrometry on B cells from 5 HD, 7 CLL, 7 FL, and 16 MCL patients (**Supplemental Table S1**). We identified 8,027 proteins in total with 1,502 proteins present in all batches (**Figure 1A and B, Supplemental Table S2**). Principal component analysis (PCA) shows that HD and individual malignancies cluster among themselves and lymphomas capable of forming solid tumors (MCL and FL) cluster similarly while CLL, which remains predominantly in the peripheral blood, clusters separately (**Figure 1C**).

**Figure 1:**
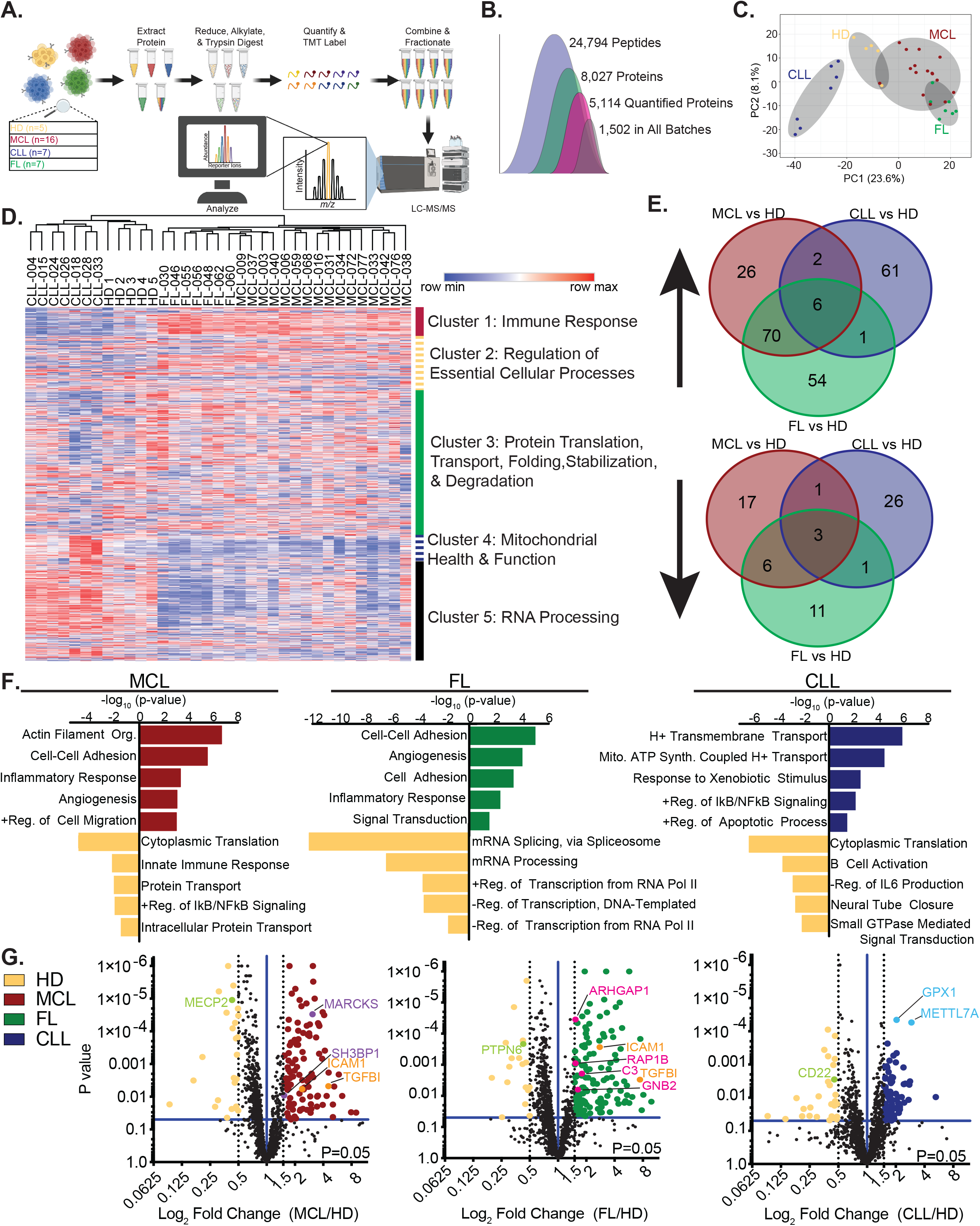
Quantitative proteomics identifies protein landscapes of lymphoid neoplasms as compared to healthy B cells. **A**. Schematic of sample processing workflow of protein from either healthy donor (HD) B cells or lymphoid tumors from three different B-cell malignancies: mantle cell lymphoma (MCL), chronic lymphoblastic leukemia (CLL), and follicular lymphoma (FL). Combined protein from all samples were processed the same as individual samples and used as a control in each run. **B**. Filtering parameters of LC-MS/MS outputs from the five different MS batches. 24,794 peptides corresponding to 8,027 proteins were filtered to only include those with an abundance output (5,114). Analysis was performed on proteins identified in all batches (1,502). **C**. PCA plot using ClustVis software of normalized abundances for all samples. **D**. Heat map of the Log_2_ value of normalized abundances for all samples. Hierarchical clustering was performed using Euclidian distance for both rows and columns. **E**. Venn Diagram of common upregulated (≥1.5-fold) (above) and downregulated (≤0.5-fold) (below) proteins between MCL, FL, and CLL all as compared to HD samples. **F**. GO terms for upregulated (≥1.5-fold) and downregulated (≤0.5-fold) proteins for each cancer as compared to HD. **G**. Volcano plots of the Log_2_ fold change for each cancer as compared to HD showing significant (p≤0.05) upregulated (≥1.5-fold) and downregulated (≤0.5-fold) proteins. Proteins upregulated in both MCL and FL depicted in orange, proteins upregulated in MCL in purple, proteins upregulated in FL in pink, proteins upregulated in CLL in light blue, and downregulated proteins of interest in light green.

Heat map of all disease protein abundances shows that samples cluster by disease sub-type and form five distinct protein clusters associated with various Gene Ontology (GO) pathways (**Figure 1D, Supplemental Tables S3-S8**). The most distinct clusters being one, four, and five. Cluster one proteins, upregulated in MCL and FL, primarily contain proteins associated with the immune response, but also contain proteins associated with cell adhesion and angiogenesis. Because these tumors can infiltrate secondary lymphoid tissues to form solid tumors, targeting these proteins may be significant for slowing disease progression. Cluster four proteins, upregulated in CLL, are associated with mitochondrial health and function. It has been reported that CLL cells are heavily reliant on mitochondrial metabolism.^12^ Therefore, proteins within this cluster may prove to be vulnerabilities within the disease ideal for targeting. Cluster five proteins are downregulated in FL and MCL and are associated with RNA processing. We have previously reported that B cells lacking the catalytic domain of the E3 ubiquitin ligase component *n*-recognin 5 (UBR5) have splicing defects.^13^ Given that UBR5 is mutated in ∼18% of MCL patients, mutations to UBR5’s catalytic domain are specific to MCL tumors, and UBR5 is the second most mutated gene in our MCL patient cohort, present in 31% of samples, proteins in this cluster may provide insight into how UBR5 mutations affect MCL pathogenesis (**Supplemental Figure S1A**).^14, 15^

As compared to HD, there are 131 significantly dysregulated proteins (104 upregulated, 27 downregulated) in MCL, 101 in CLL (70 up, 31 down), and 152 in FL (131 up, 21 down) (**Figure 1E, Supplemental Table S9**). Of the dysregulated proteins, only 6 are consistently upregulated and 3 are consistently downregulated across all three malignancies. GO analysis of dysregulated proteins in each malignancy identifies unique and similar pathways between each lymphoma type (**Figure 1F**). As discussed above, both MCL and FL have upregulation of proteins associated with cell-cell adhesion, angiogenesis, and inflammatory response. Proteins of interest include intercellular adhesion molecule 1 (ICAM1) and transforming growth factor β-induced protein (TGFBI). (**Figure 1G, orange**). High ICAM1 surface levels in FL and MCL, but low levels in CLL is linked to unfavorable patient prognosis.^16^ Although it has not been studied in MCL, transcriptional silencing of TGFBI is considered to be a potential mechanism of B-cell transformation resulting in FL.^17^ These data are in line with our proteomics data, validating our findings.

Upregulated protein pathways unique to MCL are actin filament organization and positive regulation of cell migration. They include SH3-domain binding protein 1 (SH3BP1) and myristoylated alanine-rich C-kinase substrate (MARCKS) (**Figure 1G, purple**). While upregulation of SH3BP1 is a new finding in lymphoma, it’s upregulation is associated with poor prognosis in acute myeloid leukemia (AML) patients.^18^ *MARCKS* is upregulated in MCL and has an important role in cancer spread, invasion, metastasis, and Bortezomib resistance in multiple myeloma (MM).^11^

FL has one unique upregulated pathway, signal transduction, that is associated with 19 upregulated genes. Of these genes, four of them are specific to FL: Rho GTPase-activating protein 1 (ARHGAP1), complement component c3 (C3), Ras-related protein Rap-1b (RAP1B), and guanine nucleotide-binding protein G(I)/G(S)/G(T) subunit Beta-2 (GNB2) (**Figure 1G, pink**). The most significant of these proteins, ARHGAP1, has not previously been associated with lymphoma; however, one study found that it is alternatively spliced in a TRAF6 dependent manner in myelodysplastic syndrome.^19^

Upregulated pathways in CLL are very different from MCL and FL. Some proteins associated with these pathways include glutathione peroxidase 1 (GPX1) and thiol S-methyltransferase (METTL7A) (**Figure 1G, light blue**). High GPX1 expression correlates with poor prognosis in lymphoma patients.^20^ METTL7A has not previously been associated with lymphoma but overexpression is associated with enhanced human bone marrow stem cell viability.^21^

Pathways downregulated in each malignancy are highly unique. Some potentially relevant downregulated proteins from each malignancy include methyl-CpG-binding protein 2 (MECP2) in MCL, tyrosine-protein phosphatase non-receptor type 6 protein (PTPN6) in FL, and cluster of differentiation 22 (CD22) in CLL (**Figure 1G, light green**). MECP2 interacts with the SIN3a/histone deacetylase 1/2 corepressor complex to silence BIM, a proapoptotic member of the BLC-2 family.^22^ PTPN6 is an important repressor of STAT3 activation, a transcription factor constitutively activated in a variety of cancers, including lymphoma, and is considered a tumor suppressor in AML and MM.^23^

Overall, this data identifies similarities and unique aspects of three different types of lymphoid malignancies, which have been previously unidentified or underexplored. These pathways and proteins may prove to be prognostic markers for their respective diseases and highlight potential vulnerabilities for possible treatment strategies.

### Identification of cell surfaceome

Recent success of CAR-T therapy in B-cell malignances targeting specific cell surface receptors reinforces the importance of the cell surfaceome. Of the 1,502 proteins identified in our MS analysis, 120 of them localize to the plasma membrane (**Figure 2A**). One of the defining characteristics of CLL is that it is CD5^+^, CD19^+^, and CD23^+^ with weak amounts of CD22, CD79b, and surface immunoglobulin and is FMC7^-^.^24^ FL cells are CD19^+^, CD20^+^, CD10^+^, and BCL-6^+^, but are CD5^-^ and CD23^-^.^25^ MCL cells are characterized by B-cell surface markers CD19^+^, CD20^+^, CD22^+^, CD79a^+^, CD5^+^, FMC-7^+^, and CD10^-^.^26^ Cell surface protein pattern of CD5, CD19 and CD22 was consistent with diagnostic markers (**Figure 2B**). Because CAR-T therapy is rapidly becoming the treatment of choice for patients that stop responding to other therapies, and the high incidence of relapse in lymphoid malignancies – MCL has a 100% relapse rate following initial treatments – plasma membrane proteins may prove to be new targets for CAR-T treatment. Of these 120 proteins, 48 of them are either significantly up- or downregulated in each of the three malignancies individually as compared to HD B cells (**Figure 2C**). Of these 48 proteins, 11 of them are specifically upregulated in MCL (**red text**), 8 in FL (**green text**), and 7 in CLL (**dark blue text**). 10 of these proteins overlap between MCL and FL (**orange text**), 1 overlaps between CLL and FL (**light blue text**), and two are upregulated between all three malignancies (**purple text**). We measured abundance of cell surface proteins identified in our analysis in MCL and CLL cell lines – Jeko1 and Granta519 (MCL) and HG3 (CLL) – and compared them to peripheral blood in HD (**Figure 2D**). We found that in these cell lines, some proteins, such as LGALS9, CD81, and ICAM1 matched our proteomic findings, while others such as CD5, CD22, and B2M were more variable. These differences are likely due to differences between the *in vitro* and *in vivo* microenvironments, emphasizing the importance of utilizing patient samples when identifying therapeutic targets of interest.

**Figure 2:**
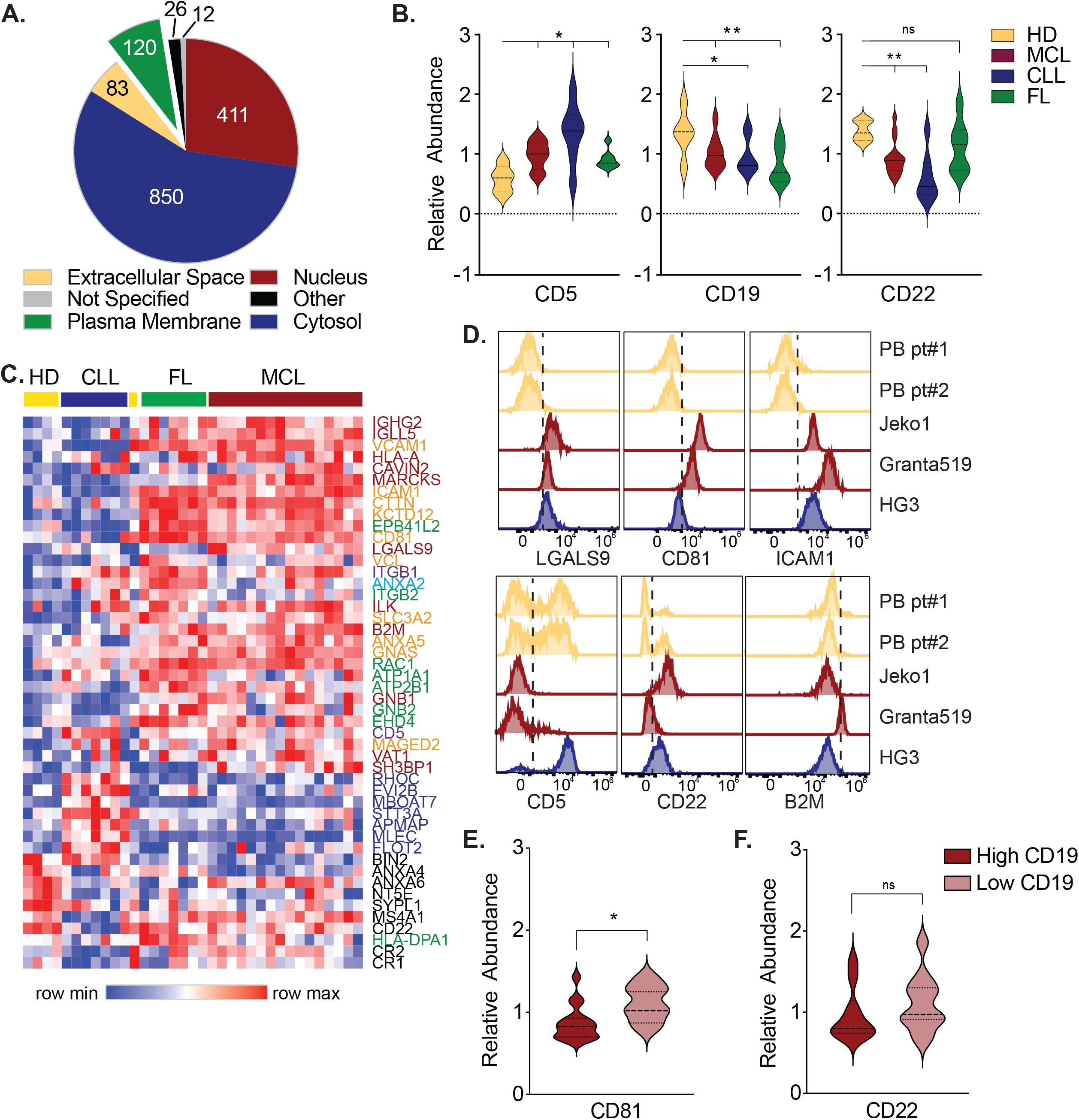
Cell surfaceome identifies potential CAR-T cell therapy targets in MCL. **A**. Pie chart of localization of 1,502 identified proteins highlighting 120 plasma membrane proteins. **B**. Relative protein abundance of cell surface markers associated with B-lymphoid malignancies (CD5, CD19, CD22). **C**. Heat map of plasma membrane proteins that are differentially expressed in at least one of the three different B lymphoid malignancies as compared to HD. **D**. Cell surface expression by flow cytometry in normal human peripheral blood (PB), and B-Lymphoid derived cell lines Jeko1 (MCL), Granta519 (MCL), and HG3 (CLL). **E**. Relative abundance of CD81 and **F**. CD22 in CD19 high and low expression patients. ns = not significant, * = p-value ≤ 0.05, ** = p-value ≤ 0.01.

Current CAR-T therapy in lymphoma targets CD19^+^ cells.^5^ It has been suggested that low CD19 may diminish efficacy of CAR-T treatment suggesting dual receptor targeting could improve outcomes.^27^ We found that CD81 was increased in MCL, but not CLL. When we stratified MCL patients based on CD19 expression, we found that CD19 and CD81 had an inverse relationship to one another, but no significant difference in CD22 expression (**Figure 2E and 2F**). This suggests that using CD81 as a CAR-T therapy target in combination with CD19 may improve patient outcomes. Together, these findings provide additional potential receptors for cellular therapy.

### Genetic heterogeneity of MCL corresponds to proteomic heterogeneity

It is thought that genetic heterogeneity is one of the reasons for the poor survival and limited treatments in MCL.^14^ The 16 MCL patients included in our study had typical age and male to female distribution. While there were mutations in 88 different genes, with patients having up to 15 mutations, we saw the expected frequency in MCL for TP53 and ATM mutations (**Supplemental Figure S1A**). PCA analysis revealed no distinct clustering in the MCL samples (**Figure 3A**). Overall mutational profile or number of mutations again revealed no specific clustering in the MCL cohort (**Supplemental Figure S1B**).

**Figure 3:**
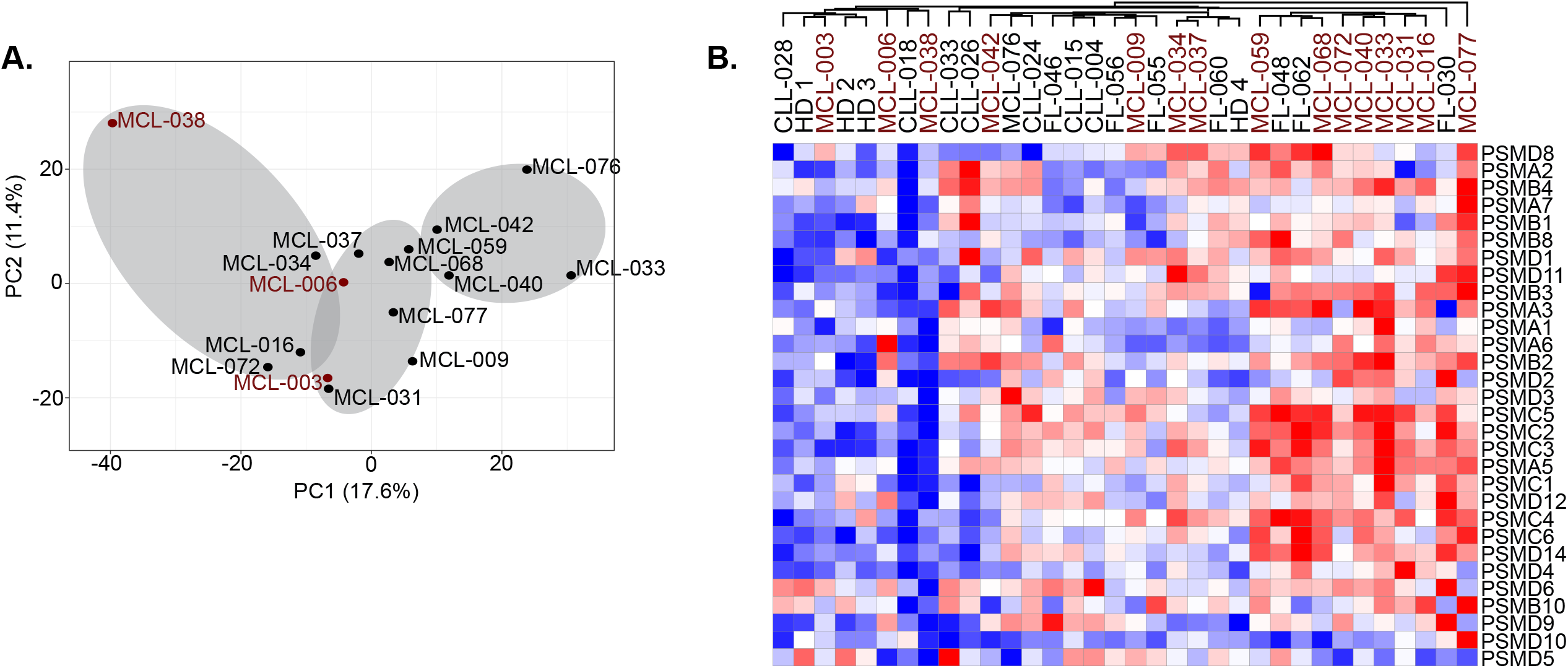
Analysis of proteasome components suggest precision targeted therapeutics could improve patient outcomes. **A**. PCA plot of MCL patient samples with clustering determined by ClustVis software. **B**. Heatmap of proteasome subunits in all patients clustered by Pearson’s Correlation.

Besides standard high dose of cytarabine regime, limited treatments are available for patients with MCL, resulting in poor survival rates and outcomes.^2^ The proteasome inhibitor Bortezomib has recently been approved for patients with relapse or refractory disease, however, there has been mixed results.^6^ It has been reported that expression of the proteasome subunits correlates with proteasome activity and increased sensitivity to proteosome inhibition.^28^ Clustering MCL, CLL, and FL patients based on protein expression of proteasome components shows diverse expression profiles among the MCL cohort (**Figure 3B**). A number of patients clustered with HD samples with low expression of proteasome components (MCL-006, MCL-003, and MCL-038) suggesting that the patients would have decreased sensitivity to Bortezomib. These findings suggest that for a heterogenous disease like MCL, precision targeted therapeutics could benefit patients and highlight the importance of using proteomics to identify new therapies that may improve patient outcomes.

### Novel post transcriptionally regulated proteins in MCL

Because MCL has the lowest 5-year survival and a high need for new therapeutics, we compared protein expression of MCL patients to both HD B cells and to the other B-cell malignancies. While there were no proteins consistently downregulated in MCL as compared to all groups, comparison of the upregulated proteins found that there are 10 proteins specifically upregulated in MCL (**Figure 4A and B, Supplemental Table S10**). These proteins are high mobility group box 3 (HMGB3), EH domain containing 1 (EHD1), MARCKS, galectin 9 (LGALS9), PDZ and LIM domain protein 1 (PDLIM1), drebrin 1 (DBN1), glutathione S-transferase Pi (GSTP1), dimethylarginine dimethylaminohydrolase 2 (DDAH2), stathmin 1 (STMN1), and four and a half LIM domains protein 1 (FHL1). Importantly, all 10 of these proteins had 3 or more unique peptides identified. (**Supplemental Table S2, Supplemental Figure S2)**. Of these, seven have never been associated with MCL and three have never been associated with any form of lymphoid malignancy (**Supplemental Figure S2**). Using previously published transcriptional data that includes these same MCL patients, we compared mRNA expression for these 10 proteins.^29^ We found that seven of them are unchanged transcriptionally (**Figure 4C**). The RNA expression level of these 10 genes shows no significant difference in patient survival, which is not surprising given that the transcripts are unchanged (**Supplemental Figure S3**).

**Figure 4:**
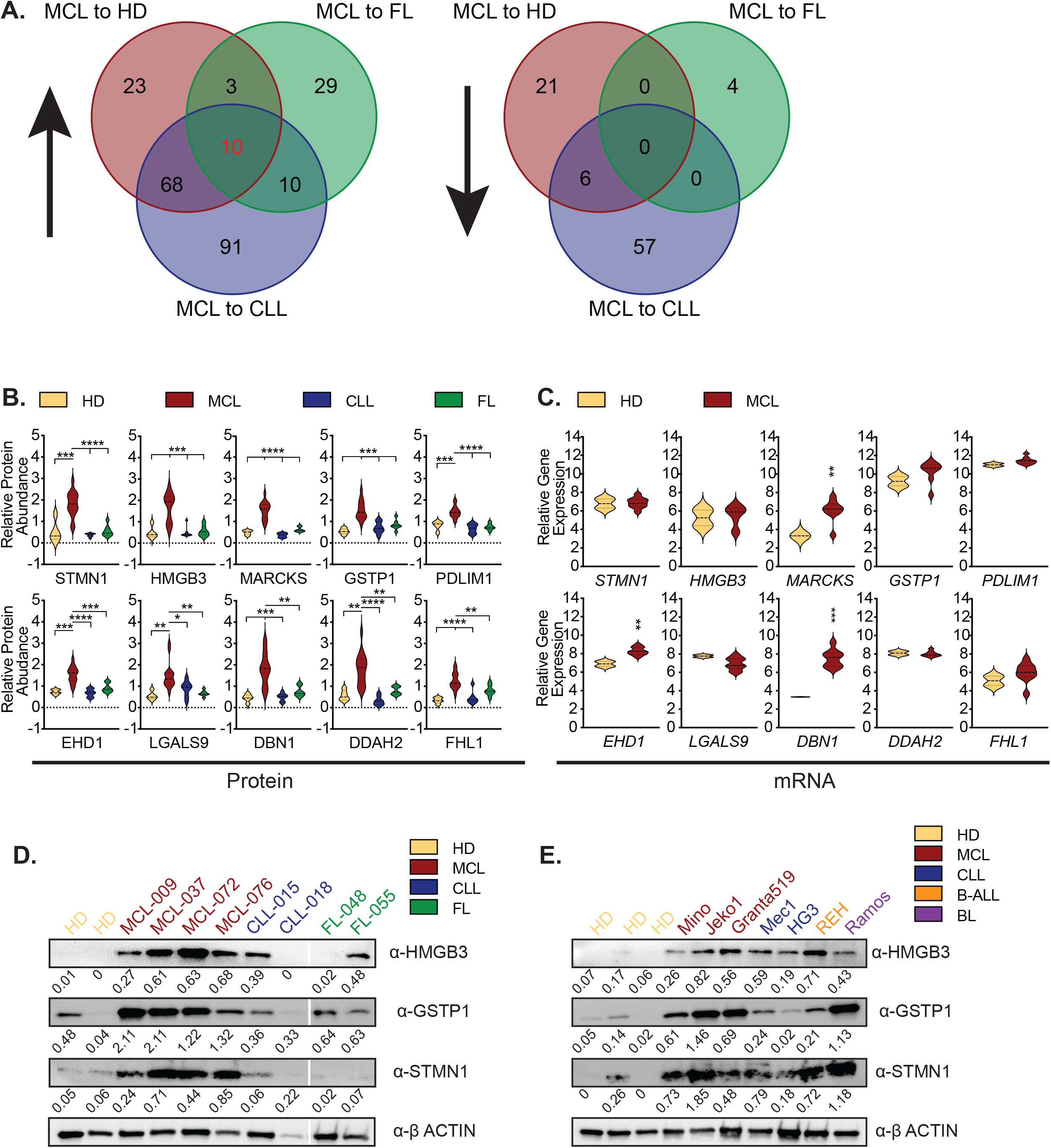
Identification of 10 post-transcriptionally regulated proteins specifically upregulated in MCL. **A**. Venn Diagram of proteins significantly (p≤0.05) upregulated (≥1.5-fold) (left) and downregulated (≤0.5-fold) (right) in MCL samples as compared to HD, FL, and CLL samples. **B**. Violin plots of relative protein abundance for 10 proteins specifically upregulated in MCL as compared to HD, CLL, and FL. **C**. Violin plots of relative gene expression as determined by RNA microarray of the 10 proteins from A in MCL as compared to HD. **D**. Western blot confirmation in select MCL, CLL, and FL primary patient samples and **E**. MCL, CLL, B-cell Acute Lymphoblastic Leukemia (B-ALL), and Burkitt’s Lymphoma (BL) patient derived cell lines as compared to HD. * = p-value ≤ 0.05, ** = p-value ≤ 0.01, *** = p-value ≤ 0.001, **** = p-value ≤ 0.0001.

To validate the upregulation of these proteins in MCL, we performed western blots for three these 10 proteins (STMN1, GSTP1, and HMGB3) in both primary patients and patient-derived cell lines including three MCL cell lines: Mino, Jeko1, and Granta519; two CLL cell lines: Mec1 and HG3; and two other lymphoma cell lines: REH, a B-cell acute lymphoblastic leukemia (B-ALL) cell line; and Ramos, a Burkitt’s lymphoma cell line (**Figure 4D and E)**. Our results in both models show these proteins are increased in MCL as compared to HD B cells and to samples from other lymphoma types.

These findings identify novel proteins that are differentially expressed at the protein level that have previously not been identified in classic transcriptional profiling, suggesting potential novel targets for drug discovery or new translational avenues for PROTAC-based drug development tailored to highly expressed proteins in MCL.

## DISCUSSION

Historically, therapeutic targets for cancer treatments have been identified using GEP. While these studies have identified many relevant targets, many of them have not been confirmed at the proteomic level. Additionally, most of these studies – including those that examine the changes to the proteome – examine only one type of cancer and compare that cancer only to HD samples. Furthermore, these analyses are heavily reliant on cell lines and when patient samples are used, the number of patients included is very small.

To gain a better understanding of the differences between hematological malignancies, we have taken samples from patients suffering from three different B-cell malignancies and utilized quantitative proteomics to identify protein signatures of each malignancy as compared to both healthy B cells and the other disease types. With the identification of over 8,000 proteins across all samples, this study provides an in-depth analysis of the proteomes of each malignancy type and utilizes the findings in MCL to validate the results. Overall, the study identifies proteins that are specifically dysregulated in MCL that may prove to be therapeutic targets for MCL therapies in the future.

Given the ever-increasing prevalence of CAR-T therapy in the treatment of B-cell malignancies, our study identified a significant number of surface proteins that may prove to be novel CAR-T targets. We also found that patients may benefit from proteomic analysis of proteasome subunits prior to Bortezomib-based treatments. One of the most striking findings of this study is the differences between the proteomic signature and the previously identified GEPs of these patients. Because most MCL patients exhibit a heterogenous mixture of genomic mutations, it is difficult to treat the disease, and this lends to the high mortality rate of these patients. We had expected to see that patients who displayed alterations in genes frequently mutated in MCL such as ATM, TP53, and UBR5 would cluster together and show similar changes in their proteome. However, we found that the number and type of mutations present in these patients had little to no effect on the proteins that are either up or downregulated. Furthermore, the finding that only three of the 10 proteins specifically upregulated in MCL also exhibited increases in mRNA reinforces the importance of examining the patient’s proteome in addition to their genome to identify therapeutic targets for treatment.

The presence of MARCKS and EHD1 in our study, two proteins whose corresponding gene expression has previously been reported to be upregulated in MCL as compared to CLL and in NHL as a whole, respectively, validates these findings.^11, 30^ In addition to MARCKS and EHD1, only one of the other 10 identified proteins had a transcriptional change, DBN1. It has previously been identified that DBN1 is a direct target of SOX11, a protein known to be aberrantly expressed in MCL.^31^ The role that these three proteins play in actin cytoskeleton formation, membrane formation, and the formation of cellular projections suggest that they may be important for disease progression and infiltration into secondary lymphoid tissues. Although it does not have a transcriptional change present like the other three, identification of GSTP1 as overexpressed at the proteomic level also validates the study. GSTP1 has been shown to be chronically overexpressed in MCL and may provide resistance to drug therapies as its inhibition increases chemotherapy sensitivity in MCL patients.^32^ Because MCL patients chronically relapse following current drug regimens, targeting GSTP1 in conjunction with these therapies may prove to be an effective strategy to improve patient outcomes.

The other six proteins specifically upregulated in MCL are also transcriptionally unchanged, indicating that they would have remained unidentified by typical patient screening methods. These six proteins have never been associated with MCL previously. STMN1 is known to be overexpressed in other types of leukemias and lymphomas.^33^ LGALS9 has been shown to be upregulated on the surface of obese B-ALL patients, and targeting this protein was reported to promote cell death, disrupt cell cycle progression, and prolong survival in B-ALL patients.34 FHL1 stabilization leads to Ibrutinib resistance in diffuse large B-cell lymphoma.35 HMGB3, PDLIM1, and DDAH2 have never been associated with any form of lymphoma previously, making them of particular interest to this study.

In conclusion, this study highlights the importance of utilizing proteomics when determining the treatment strategy for lymphoma patients. This data can be used as a resource for identifying novel treatment strategies not only in MCL, but in other B-cell malignancies. It identifies several new potential CAR-T therapy targets as well as novel proteins of interest for drug discovery or PROTAC-based drug development, highlighting the importance of the work.

## METHODS

### Patient Samples

The study was approved by the institutional review board of the University of Nebraska Medical Center (UNMC) (0161-95-EP and 0079-19-EP) and performed in accordance with the Declaration of Helsinki. Matched gene expression microarray data are available through the Gene Expression Omnibus, accession GSE132929.^14, 29^ Samples obtained from the UNMC Lymphoma Tissue Bank were prioritized for inclusion in this study if they had previously undergone pathology review, been interrogated by Affymetrix U133 Plus 2.0 gene expression microarrays, and exome sequenced.^9,28^ Patient samples are at diagnosis, and annotated for gender, age at diagnosis, OS, and mutational profile (**Supplemental Table S1**). Protein was isolated from 16 MCL, 7 FL, and 7 CLL patients. For the 5 HD patient samples, either adult CD19^+^ B cells were purchased from Stem Cell Technologies or magnetic selected B220^+^ B cells were isolated from tonsils. All samples were de-identified and accompanied by the diagnosis from the medical records, plus overall survival and patient mutations. Mutation and CNA data are publicly viewable through cBioPortal: https://www.cbioportal.org/study/summary ?id=mbn_mdacc_2013.^9^

### Mass Spectrometry

Solid tumors from MCL and FL were ground to powder in liquid nitrogen. Samples were prepared for TMT labelling, as previously described.^13^ Samples were prepared in five separate batches each containing a combined sample control (4μg from each sample) and a mixture of HD, CLL, FL, and MCL samples. Protein identification was performed using proteome discoverer software version 2.4 (ThermoFisher, Waltham, MA) by searching MS/MS data against the UniProt human protein database downloaded February 2022. Data was normalized to the average abundance of the batch control for each of the five batches. Analyses were performed on the ratio of the abundance of normalized protein in batch control to individual patient sample. Data are available via ProteomeXchange with identifier PXD038937.

### Flow Cytometry

Cells were stained for 1 h on ice with indicated antibodies in 3% FBS in PBS. Data was then analyzed using FlowJo software (Treestar, Ashland, OR). Antibody information is available in supplementary materials and methods.

### Cell Culture

Jeko-1, Mino, REH, and Ramos cells were purchased from ATCC (Manassa, VA) and Granta519, HG-3, and MEC-1 cells were purchased from DSMZ (Leibniz Institute, Germany). Cell lines were cultured following company specifications. All cells were confirmed mycoplasma negative monthly while in culture.

### Western Blot

Samples were prepared and analyzed as previously described.^13^ Antibody information is available in supplemental materials and methods.

### Statistical analysis

All experiments were performed in triplicate unless noted. Proteins with a fold change ≥ 1.5 were considered as upregulated, ≤ 0.5 as downregulated, and with a p-value ≤ 0.05 as significant. Statistical analyses were performed using unpaired two-tailed Student’s t-test assuming experimental samples of unequal variance (with Welch’s correction). Error bars depict the standard deviation ± the mean. PCA plots were generated using ClustVis online tool (https://biit.cs.ut.ee/clustvis/). Heat maps were completed using Morpheus online tool (https://software.broadinstitute.org/morpheus), using hierarchical clustering and log_2_ transformation of protein abundance. Volcano plots were made using GraphPad Prism v10.0.2 (GraphPad, La Jolla, CA). Gene ontology (GO) pathway analysis was performed using DAVID Bioinformatics Database v6.8 (https://davidbioinformatics.nih.gov/home.jsp). Outputs were first sorted by count, and then by p-value. All outputs with ≥ 5 associated genes and a p-value ≤ 0.05 are included for reference in supplementary tables. Graphs were made using the top 5 hits for both count and p-value. GO terms with similar names and gene were only listed once under the first instance. For survival curve analyses, the p-value was calculated using a Log-rank (Mantel-cox) test.

## AUTHOR CONTRIBUTIONS

S.A.S., and S.M.B. conceived and designed the experiments; S.A.S., C.B.W., K.K.D., K.J.W., and S.M.B. performed experiments and analysis, H.C.H.L., and N.T.W. provided technical and material support; J.M.V., T.G., and M.R.G. provided samples and/or clinical data. S.A.S. and S.M.B. wrote the manuscript. All authors reviewed the manuscript before submission.

## ACKNOWLEDGEMENTS

We would like to thank the University of Utah Flow Cytometry Shared Resource Laboratory, and UNMC Mass Spectrometry and Proteomics Core Facility for expert assistance. The Nebraska Lymphoma Tissue Bank and core facilities are administrated through the Office of the Vice Chancellor for Research and supported by state funds from the Nebraska Research Initiative (NRI) and The Fred and Pamela Buffett Cancer Center’s National Cancer Institute Cancer Support Grant (P30CA036727). S.M.B. is supported by the National Institutes of Health P20GM121316, R37CA262635, and R01AI53090. This publication was supported by the Huntsman Cancer Institute at the University of Utah, supported by the National Cancer Institute of the National Institute of Health under award number P30CA042014.

