## Supplemental Materials and Methods for "Proteome landscape of B-cell malignancies identifies mantle cell lymphoma protein signature"

#### Flow Cytometry Antibodies:

| Antibody | Fluorochrome | Ref# | Clone | Company |
|---|---|---|---|---|
| LGALS9 | FITC | 348911 | 9M1-3 | BioLegend |
| CD81 | FITC | 349503 | 5A6 | BioLegend |
| ICAM1 | PacBlue | 322715 | HCD54 | BioLegend |
| CD5 | APC | 364015 | L17F12 | BioLegend |
| CD22 | APC | 363505 | S-HCL-1 | BioLegend |
| B2M | PE | 316305 | 2M2 | BioLegend |

#### Western Blot Antibodies

| Antibody | Company | Ref# | Dilution |
|---|---|---|---|
| HMG2a (HMGB3) | Bethyl | a300-736a-t | 1:1000 |
| GSTP1 | Proteintech | 15902-1-ap | 1:1000 |
| STMN1 | Proteintech | 11157-1-AP | 1:1000 |
| β-ACTIN-HRP | Santa Cruz | Sc-47778 | 1:10000 |

### SUPPLEMENTAL FIGURE LEGENDS

#### Supplemental Figure 1

**A.** Table indicating mutations present in each mantle cell lymphoma (MCL) sample with the frequency of that mutation in all MCL samples by percentage (%). **B.** Heat map of all proteins identified in MCL patients clustered by Pearson's Correlation.

#### Supplemental Figure 2

**A.** 10 proteins specifically upregulated in MCL with their number of unique peptides, fold change and p-value in MCL as compared to HD, protein function, role in lymphoma as a whole, and role in MCL specifically.

#### Supplemental Figure 3

**A.** Survival curves of MCL patients based on publicly available gene expression data for the 10 proteins specifically upregulated in MCL.[29]
