## Supplementary figures and images for "Proteome landscape of B-cell malignancies identifies mantle cell lymphoma protein signature"

### Supplemental Figure 1

**Swenson, et al. MCL Patients Supplemental Figure S1**

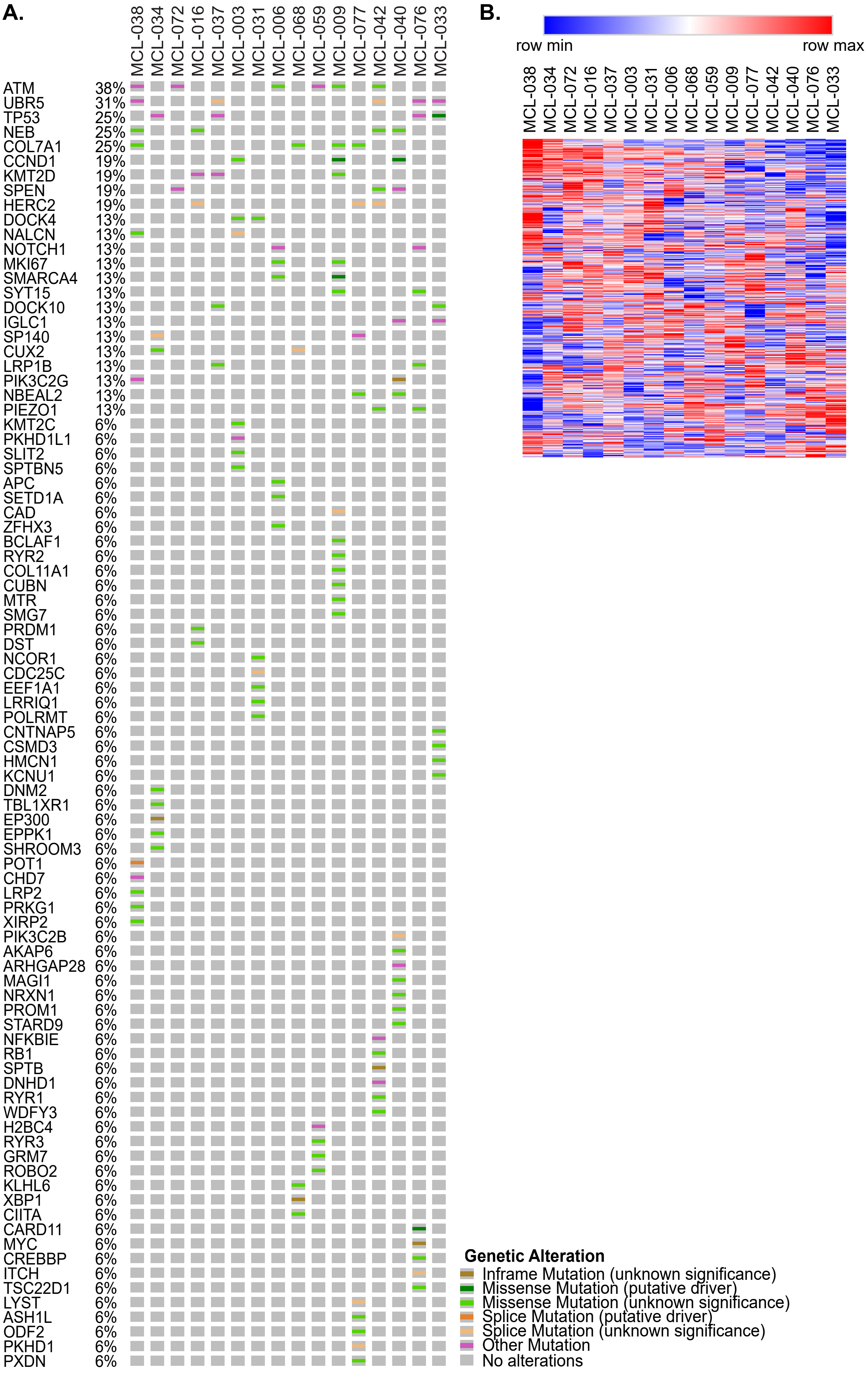

### Supplemental Figure 3

**Swenson, et al. MCL Patients Supplemental Figure S3**

**A.**

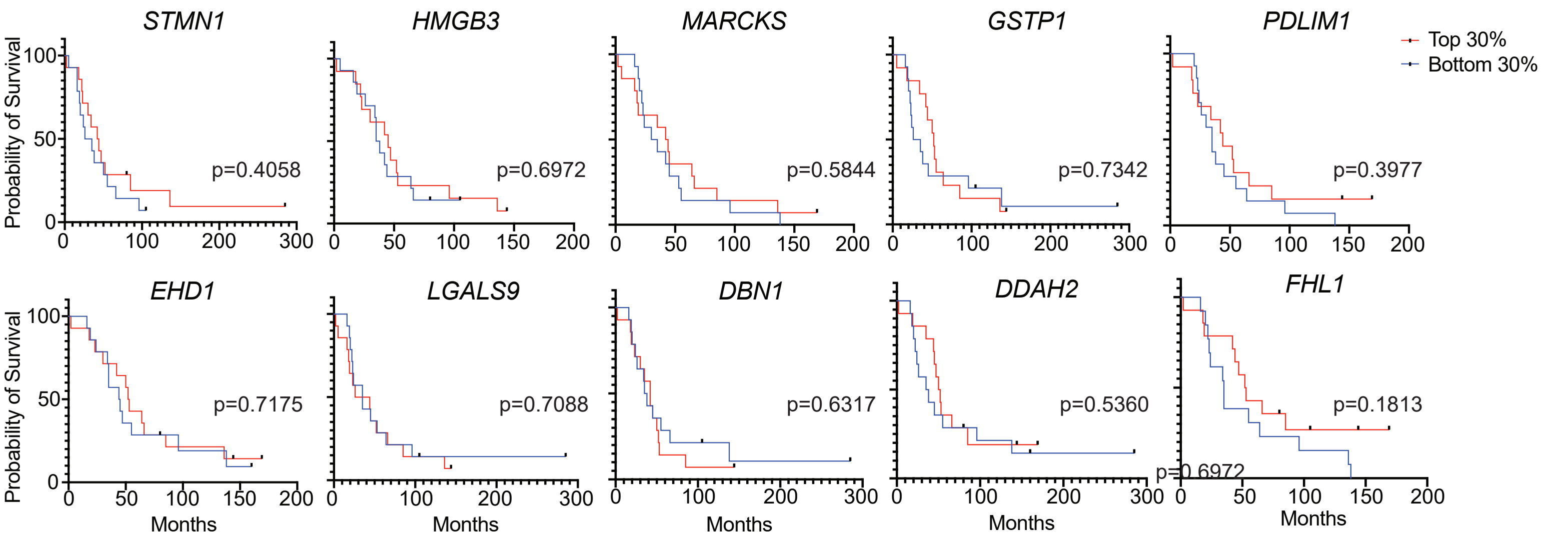
