## Supplemental Figure 2 for "Proteome landscape of B-cell malignancies identifies mantle cell lymphoma protein signature"

**Swenson, et al. MCL Patients Supplemental Figure S2**

| Protein Name & Symbol | #Unique Peptides | MCL/HD Protein Abundance & p-value | Function | Role in Lymphoma | Role in MCL |
|---|---|---|---|---|---|
| Drebin 1 (DBN1) | 4 | 4.01 8.8E-04 | Actin cytoskeleton-organizing protein, assists with forming cell projections and recruiting CXCR4 to immunological synapses | Overexpressed in ALL[31] | Direct targets of SOX11 in MCL[31] |
| Four and a Half LIM Domains Protein 1 (FHL1) | 6 | 3.64 5.9E-05 | Plays a crucial role in assembly of sarcomeres, signal transduction, and muscle growth | Stabilized FHL1 leads to Ibrutinib resistance in Diffuse Large B-Cell Lymphoma[35] | None known |
| High Mobility Group Box 3 (HMGB3) | 4 | 3.5 9.5E-04 | DNA binding, Negative Regulator of B and Myeloid cell differentiation | None known | None known |
| Stathmin 1 (STMN1) | 3 | 3.498 2.5E-04 | Microtubule destabilization | Overexpressed in many leukemias and lymphomas[33] | None known |
| Dimethylarginine Dimethylamino-hydrolase 2 (DDAH2) | 6 | 3.2 1.7E-03 | Regulates nitric oxide bioavailability | None known | None known |
| Myristoylated Alanine Rich Protein Kinase C Substrate (MARCKS) | 6 | 3.13 1.8E-05 | Actin cross-linking protein | Targeting p-MARCKS overcomes drug resistance in multiple myeloma[11] | Transcriptionally upregulated in MCL vs CLL[11] |
| Glutathione S-Transferase pi 1 (GSTP1) | 8 | 2.83 1.2E-04 | Conjugates reduced glutathione to hydrophobic electrophiles | Allows for detoxification of cytotoxic drugs making drug treatments difficult and relapse common[32] | Chronically overexpressed, inhibition increases sensitivity to chemotherapy[32] |
| Galectin 9 (LGALS9) | 3 | 2.67 3.5E-03 | Binds galactosides | Upregulated in B-Acute Lymphoblastic Leukemia (ALL) and antibody-mediated targeted results in increased survival[34] | None known |
| EH Domain Containing 1 (EHD1) | 14 | 2.12 1.6E-04 | Regulates trafficking of receptors and cargo from the endocytic recycling compartment | Upregulated in Non Hodgkin Lymphoma[20] | None known |
| PDZ and LIM Domain 1 (PDLIM1) | 11 | 1.76 1.8E-04 | Involved in assembly/ dissassembly and directioning of stress fibers in fibroblasts | None known | None known |
